# Building modern coexistence theory from the ground up: the role of community assembly

**DOI:** 10.1101/2023.01.13.523886

**Authors:** Jurg W. Spaak, Sebastian J. Schreiber

**Author notes:** Corresponding author Email addresses.

## Abstract

Modern coexistence theory (MCT) is one of the leading methods to understand species coexistence. It uses invasion growth rates – the average, per-capita growth rate of a rare species – to identify when and why species coexist. Despite significant advances in dissecting coexistence mechanisms when coexistence occurs, MCT relies on a “mutual invasibility” condition designed for two species communities, but poorly defined for species rich communities. Here, we review well-known issues with this component of MCT and propose a solution based on recent mathematical advances. We propose a clear framework for expanding MCT to species rich communities and for understanding invasion resistance as well as coexistence, especially for communities which could not be analyzed with MCT so far. Using two data-driven community models from the literature, we illustrate the utility of our framework and highlight the opportunities for bridging the fields of community assembly and species coexistence.

*Statement of authorship*: Studied conceived jointly by JWS and SJS. JWS and SJS wrote the manuscript together. JWS wrote the python code and SJS wrote R code.

*Data accessibility*: All computer code used in this manuscript will be made publicly available on figshare.

Niche and fitness differences | Storage effect | Coexistence

## Introduction

The richness and composition of co-occurring species can exhibit dramatic variation across space and time. Understanding the processes generating and maintaining this variation is a central focus of community ecology (Chase, 2003; HilleRisLambers *et al*., 2012; Mittelbach & Schemske, 2015). Dispersal of individuals from a regional species and filtering by the abiotic environmental conditions determines which species have the opportunity to co-occur locally (Kraft *et al*., 2015a). As populations grow, species interactions and dynamics determine which subsets of species coexist and which of these subsets resist invasion attempts from the rest of the regional species pool (Law & Morton, 1996a; Chase, 2003; HilleRisLambers *et al*., 2012). Hence, understanding what factors determine persistent patterns of species co-occurrence requires understanding the process of community assembly, coexistence mechanisms, and the determinants of competitive exclusion, or more generally invasion resistance.

At an ecological time scale, three outcomes of community assembly are possible (Law & Morton, 1996a; Chase, 2003). First, regardless of the order in which the species invade, the community always assembles to the same configuration of species – a unique stable state. Second, there are alternative stable states and community assembles to one of these states but which one may depend on the order which species arrive to the local site. Finally, community assembly may never settle. There is a constant cycling in species composition as invasions from the species pool repeatedly shift the community from one state to another state. Despite theoretical support for cycling (Yodzis, 1978; Schreiber & Rittenhouse, 2004; Allesina & Levine, 2011), it is rarely observed in empirical studies (Chase, 2003; Warren *et al*., 2003).

Many mechanisms have been proposed to explain why some collections of species coexist, while others do not (Chesson, 2000). These mechanisms include competition for limiting resources (Tilman *et al*., 1982; MacArthur, 1970; Letten *et al*., 2017), shared predators (Paine, 1966; Janzen, 1970; Holt & Lawton, 1994) or mutualists (Johnson & Bronstein, 2019), and temporal fluctuations in environmental conditions (Chesson, 1994, 2003). Modern coexistence theory (MCT) attempts to unify our understanding of these different concepts by disentangling mechanisms into their stabilizing and equalizing components (Chesson, 2000).

The key ingredient of MCT is the per-capita growth rate of a species (the invader) at low density while other species (the resident community) are at equilibrium or exhibit long-term stable fluctuations– the invasion growth rate (MacArthur & Levins, 1967; Turelli, 1978; Chesson, 2000; Schreiber, 2000; Grainger *et al*., 2019b). A positive invasion growth rate implies that the invader tends to increase when rare, while a negative invasion growth rate implies that the resident community resists the invasion attempt. MCT is based on the assumption that if the “right” invasion growth rates are positive then coexistence occurs. MCT, then, decomposes invasion growth rates into different coexistence mechanisms to identify their relative importance (Chesson, 2003, 1994; Ellner *et al*., 2019; Chesson, 2020; Spaak *et al*., 2021b).

But what are the “right” invasion growth rates and in what sense does their being positive ensure coexistence? The answer is clear for two species competitive communities. Each species should have a positive invasion growth rate when the other species is at equilibrium, i.e the mutual invasability criterion. But what about more complex communities? The most commonly taken approach is to test whether each species has a positive invasion growth rate while it is absent and the resident community is at equilibrium. Defined as such, these invasion growth rates may not exist or may not be unique (Spaak *et al*., 2021a; Barabás *et al*., 2018; Saavedra *et al*., 2017). Additionally, even when they are defined, positive invasion growth rates might not be sufficient to guarantee coexistence; an important exception, however, are competitive communities with diffusive interactions–comparable interaction strengths for all species pairs (Chesson, 2018). This is especially unfortunate, as MCT explores more and more species rich communities with asymmetric interactions (Spaak *et al*., 2021a; Shoemaker *et al*., 2019; Godoy *et al*., 2018; Saavedra *et al*., 2017; Petry *et al*., 2018; Chesson, 2018; Spaak & De Laender, 2021). Knowledge about these two issues are wide-spread in MCT (Spaak *et al*., 2021b; Spaak & De Laender, 2020; Barabás *et al*., 2018; Saavedra *et al*., 2017; Pande *et al*., 2020; Chesson, 2020, 2018), yet the invasion growth rates remain widely used as if these problems did not exist (Grainger *et al*., 2019a; Letten *et al*., 2018; Shoemaker *et al*., 2019; Schreiber *et al*., 2018; Buche *et al*., 2022; Ellner *et al*., 2019; Spaak *et al*., 2022a). A

Invasion growth rates also determine whether a configuration of coexisting species is invasion resistant (Chesson & Ellner, 1989; Schreiber, 2000; Benäım & Schreiber, 2019). If the invasion growth rates are negative for all missing species, then the local community is invasion resistant and an end state for community assembly. In principle, one could use the methods of MCT to partition invasion growth rates and identify the mechanisms underlying invasion resistance. For example, to what extent is invasion resistance due to an environmental filter or having too much niche overlap with the resident species? However, we are not aware of any study explicitly performing such an analysis for a species rich community. This is surprising as in several MCT studies only a subset of the species are predicted to coexist by the models and the methods of MCT are only applied to these subsets of species (Godoy *et al*., 2014; Letten *et al*., 2018; Maynard *et al*., 2019; Spaak *et al*., 2021a). However, this misses the opportunity to understand why the models are predicting these subcommunities are invasion resistant.

Here, we build a bridge between community assembly theory and MCT to address these limitations. This bridge is based on a recent advance in the mathematical theory of permanence (Hofbauer & Schreiber, 2022) that capitalizes on earlier, more abstract theory (Schreiber, 2000; Hofbauer & Schreiber, 2010; Roth *et al*., 2017; Patel & Schreiber, 2018a). In short, this theory associates invasion growth rates with all sub-community dynamics to describe all possible transitions between communities due to single or multiple species transitions – the invasion graph. Using these graphs, one can identify, in a mathematically rigorous manner, the end states of community assembly i.e. subsets of species that coexist and are invasion resistant. For these end states, we propose to apply the MCT approach to (i) the invasion growth rates of coexisting species to understand why the species coexist and (ii) and the invasion growth rates of the excluded species to understand why the community is invasion resistant. We apply our approach to two datasets from the literature which previously could not be analyzed by MCT.

Our paper consists of four sections which can be read partially independent of each other:

1. Review how invasion growth rates are typically used in modern coexistence theory and its issues.
2. Review a mathematically rigorous treatment of invasion growth rates (i.e. permanence theory) and show how this improves the current state.
3. Propose an extension of modern coexistence theory based on permanence theory, which allows the analysis of certain species rich communities currently outside of the grasp of modern coexistence theory.
4. Apply the newly developed theory to two communities from the literature showcasing how we get new insights using these advances. We have one application using niche and fitness differences based on Lotka-Volterra dynamics as well as one application to a community with a fluctuating environment with storage effects and relative non-linearities.

We attempted to write (1) and (2) in way accessible to readers without a strong mathematical background, to do so we moved some of the mathematical subtleties and precision to Appendix A and refer to Hofbauer & Schreiber (2022) for a more precise treatment. Importantly, we encourage readers without a deep understanding of (1) and (2) to continue with the extension in (3) and applications in (4) that require less mathematical sophistication. Additionally, we provide automated code to help with empirical applications of these new concepts.

## The naïve invasion criterion

To describe the framework, we focus on continuous-time models of *n* interacting species with densities *N* = (*N*_1_, *N*_2_, …, *N_n_*). To allow for population structure (e.g. discrete habitat patches, stages, genotypes), temporal forcing (e.g. periodic or chaotic environmental fluctuations), and environmental feedbacks (e.g. plant-soil feedbacks), we allow for a finite number *m* of auxiliary variables *A* = (*A*_1_, *A*_2_, …, *A_m_*), as proposed by Patel & Schreiber (2018b) and extended to stochastic models by Benaïm & Schreiber (2019).

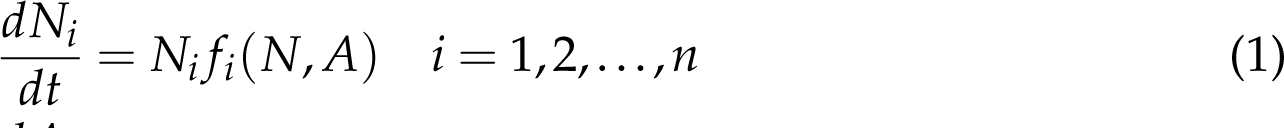

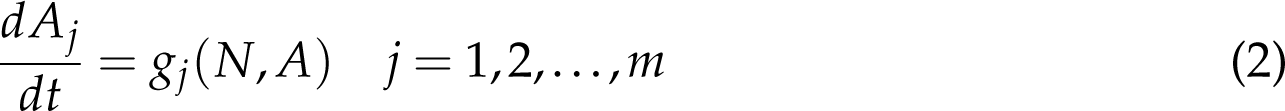

where *f_i_*is the per-capita growth rate of species *i* and *g_j_*governs the dynamics of the *j*-th auxiliary variable. There are multiple definitions in the theoretical literature about what coexistence means for these models (Schreiber, 2006). For invasion-based metrics, we argue that the concept of permanence for deterministic models (stochastic persistence for stochastic models) is most appropriate (Schuster *et al*., 1979; Sigmund & Schuster, 1984; Butler & Waltman, 1986; Schreiber *et al*., 2011). Permanence corresponds to a global attractor bounded away from extinction, i.e. after an initial transient phase, all species have positive densities above a certain threshold, even after rare large disturbances (e.g. disturbances that reduce species densities by a high percentage). Unlike classical equilibrium based concepts, permanence allows for coexistence with equilibrium or non-equilibrium dynamics (e.g. periodic, chaotic dynamics or stochastic fluctuations, see appendix Appendix D). However, consistent with limitations of invasion growth rate approaches, a system may have a stable equilibrium with all species present yet not be permanent due to the presence of an alternative stable state supporting only a subset of species (e.g. Fig. 2 D, E).

Mathematical theory for characterizing permanence or stochastic persistence using invasion growth rates has been developing for several decades (Hofbauer, 1981; Chesson, 1982; Chesson & Ellner, 1989; Ellner, 1989; Schreiber, 2000; Hofbauer & Schreiber, 2010; Schreiber *et al*., 2011; Patel & Schreiber, 2018b; Hening & Nguyen, 2018; Benaïm & Schreiber, 2019; Hening & Nguyen, 2018; Hening *et al*., 2021, 2022).

Invasion growth rates, here, correspond to the average per-capita growth rate of species missing from a subcommunity of the *n* species. Hence, an invasion growth rate, in general, depends on the sub-community context. By context, we mean some long-term stationary behavior of the model for a subset of species *S*. This stationary behavior may correspond to equilibrium or non-equilibrium dynamics (see appendix Appendix D for non-equilibrium dynamics). In the case of an equilibrium, (*N̂*, *Â*), the invasion growth rate of a species *i* absent from this equilibrium (i.e. *N̂_i_* = 0) is simply its per-capita growth rate at the equilibrium

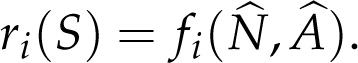

When this invasion growth rate is positive, species *i* is predicted to increase at an exponential rate when introduced at low densities into the community. When *r_i_*(*S*) is negative, species *i* is predicted to decrease at an exponential rate.

MCT rests on the principle that positive invasion growth rates imply coexistence (Grainger *et al*., 2019b; Ellner *et al*., 2019; Spaak & De Laender, 2020; Barabás *et al*., 2018). A commonly used version of this principle, what we call the naïve invasion criterion, goes back to MacArthur & Levins (1967) who wrote “such a com-munity can retain all *n* species if any one of them can increase when rare, i.e., when [species *i*’s density] *X_i_* is near zero and all the others are at the equilibrium values [or stationarity] which they would reach in the absence of *X_i_*.” From the start this was merely a heuristic criterion and did not actually guarantee coexistence (Turelli, 1978, 1981; Chesson & Ellner, 1989). There are three conceptual issues with this heuristic. First, not all of the *n* − 1 subcommunities may exist and the invasion analysis cannot be performed. A typical example is a two species predator prey community in which the 1 species subcommunity consisting only of the predator can not persist without its prey. In empirical applications of MCT, this limitation has caused most problems (Saavedra *et al*., 2017; Spaak *et al*., 2021a; McPeek, 2022). Second, all invasion growth rates might be positive for all possible *n* − 1 subcommunities, yet the species may not actually coexist (Wang *et al*., 2011). This can arise when some species experience an Allee effect (Gil *et al*., 2019; Wang *et al*., 2011). For example, Gil *et al*. (2019) considered competing species using intra- and inter-specific social information to reduce predation risk. When predation risk is high and interspecific social information substantially lowers this risk, invasion growth rates at the single species equilibria can be positive despite predation driving both species extinct whenever they are simultaneously at low densities (Fig. 2D). Third, this heuristic doesn’t provide a way to deal with only a subset of the species coexisting. Community assembly, often, may lead to an end state with only a subset of the available species. Yet, we should be interested in why aren’t the other species able to invade?

### A review of permanence theory

To address the issues associated with the naïve invasion criteria, we review the work of Hofbauer & Schreiber (2022) who introduced the concepts of an invasion scheme and an invasion graph. The invasion scheme corresponds to the invasion growth rates at all possible sub-communities, not only the *n* − 1 subcommunities. Specifically, for any sub-community *S* supporting an equilibrium (*N̂*, *Â*) (or, more generally a stationary distribution *m_S_*(*N̂*, *Â*)*dNdA* – see appendix Appendix D), we compute the invasion growth rates *r_i_*(*S*) for all the species (note that *r_i_*(*S*) = 0 for all *i* in *S* by definition). The invasion scheme is the matrix *r_i_*(*S*) with the subcommunities *S* as the rows and the species as the columns (Fig. 1).

**Figure 1:**
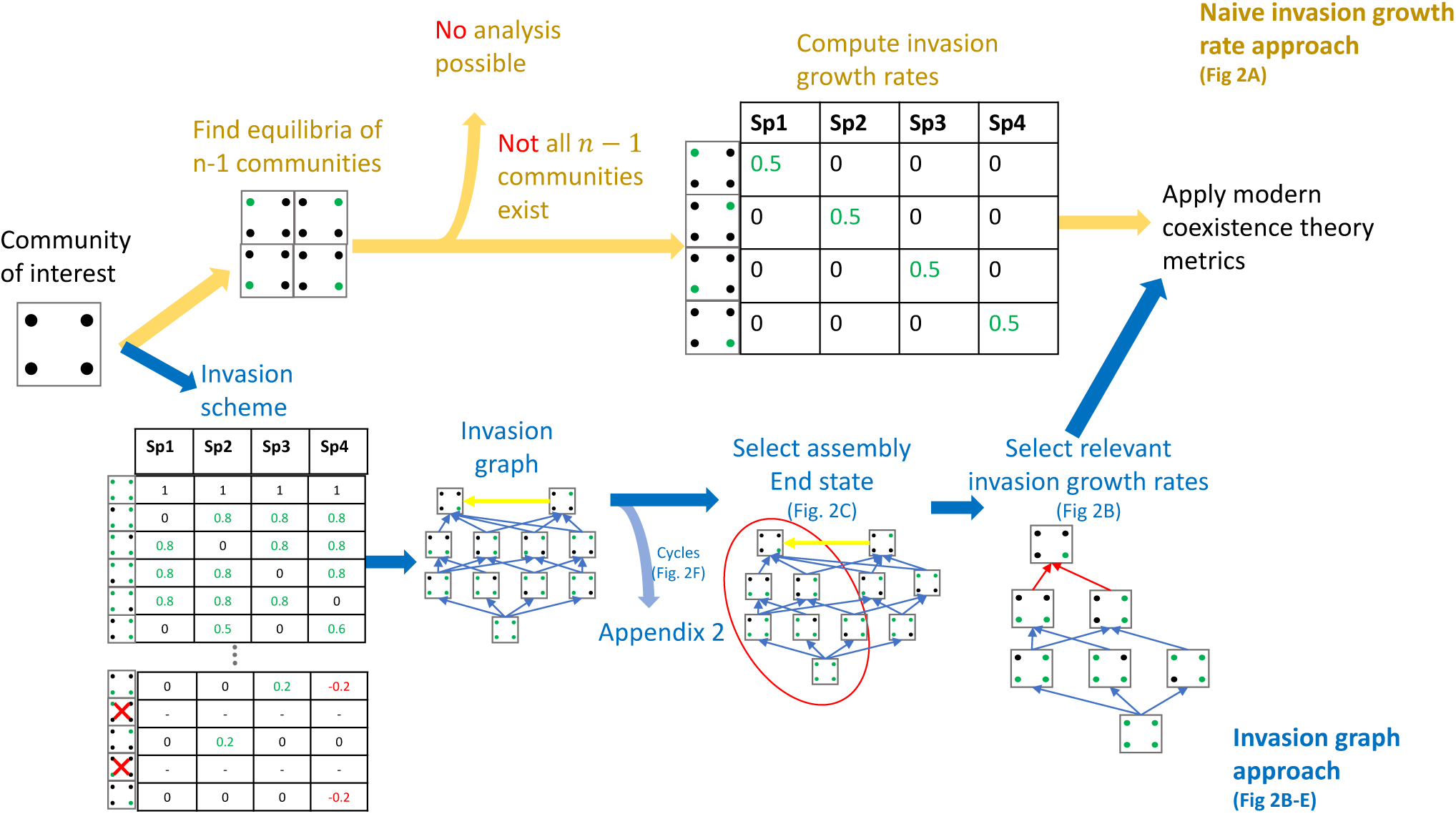
We propose a new work flow for modern coexistence theory (MCT). For a given community of interest, the typical workflow of MCT (yellow arrows) first computes the invasion growth rate of each species into the respective resident community. Then, if all *n* 1 communities, and hence invasion growth rates, exist one can apply the metrics of MCT. If not all *n* 1 communities exist no analysis is possible. Instead (blue arrows), we propose to compute the entire invasion scheme, i.e. identify equilibria (or stationary distributions) of all possible subcommunities and compute their invasion growth rates. Given this invasion scheme, we compute the invasion graph. If this invasion graph is acyclic, we select an end-state for which all species coexist and no other species could invade (red circle). Finally, we apply the invasion growth rate partitioning methods of MCT to the positive invasion growth rates associated with all realized *n* 1 subcommunities and the negative invasion growth rates for all species missing from the community (red arrows).

Next we define the invasion graph (Fig. 2), which is related to other graphs from community assembly theory (Law & Blackford, 1992; Law & Morton, 1996a; Morton *et al*., 1996; Lockwood *et al*., 1997; Song *et al*., 2021; Serván & Allesina, 2021), but is defined purely in terms on the invasion growth rates. The subcommunities *S* are the vertices of the graph and the edges between vertices are potential transitions due to invasions. Specifically, there is a directed edge from *S* to *T* ≠ *S* if (i) all species *i* in *T* but not in *S* could invade the subcommunity *S*, i.e. *r_i_*(*S*) *>* 0, and (ii) all species *j* in *S* but not in *T* could not invade the subcommunity *T*, i.e. *r_j_*(*T*) *<* 0. Intuitively, the first condition implies that all species gained in the transition from *S* to *T* can indeed invade subcommunity *S*. The second condition implies that all species lost in the transition are properly lost and could not re-invade *T*. The relevance of these transitions to the community dynamics follow from a key result of Hofbauer & Schreiber (2022): if there is a community trajectory (*N*_1_(*t*),…, *N_n_*(*t*), *A*_1_(*t*),…, *A_m_*(*t*)) that in forward time converges to an equilibrium supporting the species in *T* and that in backward time converges to an equilibrium supporting the species in *S*, then the invasion graph includes the transition *S* → *T* (see Appendix A for more details). Understanding all possible ways that community trajectories can connect community equilibria (or stationary distributions) is essential for the mathematical theory. Importantly, a transition *S* → *T* may involve species outside of subcommunities *S* and *T*. For example, Figure 2E shows an invasion graph where the transition from a subcommunity consisting of 3 coexisting competitors (*S* = {1, 2, 3}) to a sub-community consisting of one competitors (*T* = {1}) occurs due to the invasion of a predator (4).

**Figure 2:**
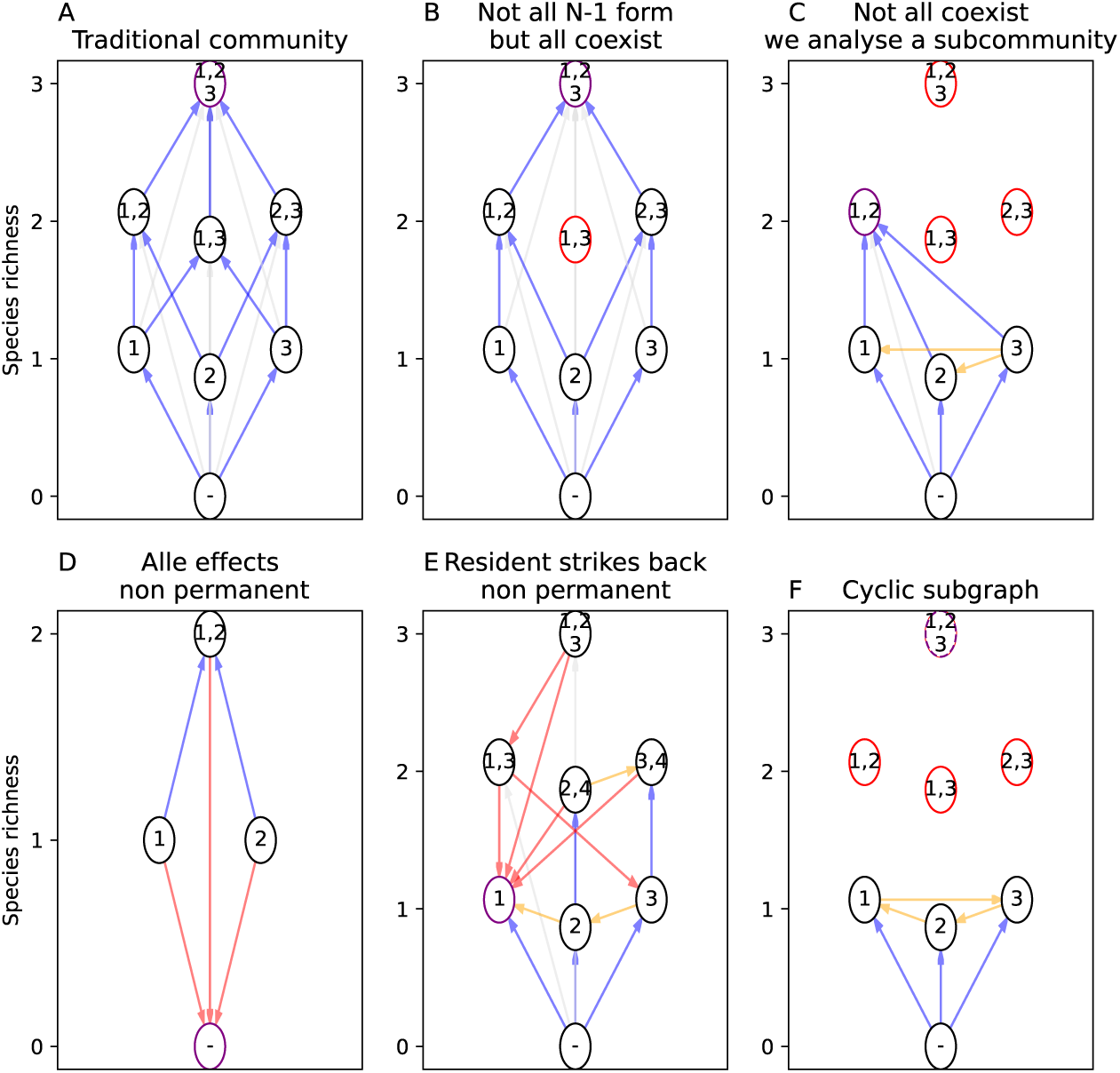
6 examples of invasion graphs corresponding to a community; A: where all missing species have positive invasion growth rates and subcommunities exist; B: where not all *n* 1 = 2 subcommunities can coexist, yet the full community can coexist; C: where not all species can coexist, but there is unique end state 1, 2. D: with Allee effects with strong inter- and intra-specific facilitation, i.e. the subcommunities are all feasible and stable, but upon loss of a species this species can not be reintroduced, such a community is not permanent. E: with a resident strikes back phenomena. Species 4 (a predator of species 2 and 3) can invade community 1, 2, 3 resulting in species 1 competitively excluding species 2 and 3. This results in species 4 going extinct, despite its positive invasion growth rate. F: with an invasion graph with a cycle (1 3 2 1). Consequently, species coexistence depends on more than the signs of the invasion growth rates. Traditional MCT only applies to the community in panel A, while the new method applies to communities A-E. The black and purple circles are realized subcommunities and height corresponds to the species richness of the subcommunity. Red circles correspond to non-realized subcommunities, purple circles correspond to potential end states. The directed edges show the transitions from one subcommunity to another due invasions (see text). Blue arrows indicate transitions which increase species richness by one, grey arrows indicate transitions increasing species richness by more than one, yellow arrows indicate transitions with no change in species richness and red (see Fig. 3) arrows indicate transitions which decrease species richness.

The invasion graph can be viewed as a representation of community assembly under three key assumptions: (i) there is a fixed species pool (i.e. the species included in equation (5)), (ii) invasions involve (infinitesimally) small propagules, and (iii) invasions are sufficiently separated in time that the species’ densities and the auxiliary variables reach their new equilibrium state (or stationary distribution). These assumptions are consistent with classical, community assembly theory (Post & Pimm, 1983; Law & Blackford, 1992; Law & Morton, 1996a; Morton *et al*., 1996; Lockwood *et al*., 1997; Serván & Allesina, 2021). While all transitions in the invasion graph are critical for evaluating coexistence, one may consider only a subset of these transitions for a model of community assembly. For example, classical assembly theory usually focuses on single species, invading propagules (see, however, the “1066 effect” in Lockwood *et al*. (1997) due to multispecies invasions).

An invasion graph can either contain a cycle or not. A cycle is a sequence of invasions starting at a subcommunity *A* eventually returns to this subcommunity e.g. invasions cause a transition from the subcommunity *S*_1_ = *A* to subcommunity *S*_2_, from subcommunity *S*_2_ to subcommunity *S*_3_,…, and ultimately from subcommunity *S_m_*back to subcommunity *S*_1_. The classic example of directed cycle is the rock-paper-scissor community (May & Leonard, 1975) where the cycle is of length *m* = 3 and the subcommunities consist of single species (Fig. 2F). More complex cycles arise naturally in multi-trophic communities (Law & Morton, 1996a; Schreiber & Rittenhouse, 2004; Spaak *et al*., 2023). When there are no directed cycles, the invasion graph is acyclic. We focus on these acyclic graphs as they appear to be more common in empirical studies (Chase, 2003) and, more to the point, the mathematical theory can only deal with certain types of cyclic invasion graphs (see Appendix B) and MCT has yet to develop an approach that applies to cyclic invasion graphs. Notably, unlike acyclic invasion graphs, cyclic invasion graphs may not identify key transitions associated with invasions. For example, even if the rock-paper-scissor community coexists, the invasion graph doesn’t identify the transition from the single species to the three species communities. This failure stems from communities trajectories that converge in forward time to the three species equilibrium do not converge in backward time to any single species equilibrium.

For acyclic invasion graphs, Hofbauer & Schreiber (2022) proved that the commu-nity is permanent if and only if for each of the subcommunities *S* there is at least one species *i* which can invade, i.e. *r_i_*(*S*) *>* 0. Importantly, these are not heuristics, such as the naïve invasion criterion, but rather mathematically rigorous results under suitable technical assumptions (Hofbauer & Schreiber, 2022) – see Appendix A for a discussion of these assumptions. Importantly, this invasion graph solves all the previous issues of the naïve invasion criterion.

The first issue of naïve invasion growth rates, that not all *n* − 1 subcommunities might exist causes no problem. The invasion graph is defined for all subcommunities *S* which do exist, if some do not exist they are simply not part of the invasion graph (Fig. 2B). Communities with Allee effects are frequent examples for the second issue, i.e. all naïve invasion growth rates might be positive yet the community does not coexist (Barabás *et al*., 2018; Schreiber *et al*., 2019). In these examples the invasion growth rates of the invader into the *n* − 1 sub-communities might be positive, but the invasion growth rates into the empty state, i.e. *S* being the empty set, is not positive (Fig. 2D). This implies that the species cannot coexist in the sense of permanence. It is possible, however, that there are alternative stable states including one supporting all species and another corresponding to extinction of all species. Hence, coexistence may be possible in a weaker sense than permanence. For the third issue, the invasion graph identifies which subcommunities are community assembly end points (Fig. 2C,E). These are subcommunities *S* such that (i) the invasion graph restricted to the species in *S* ensures coexistence, and (ii) the invasion growth rates *r_i_*(*S*) of all species *i* not in *S* are negative. For an acyclic invasion graph there is at least one end state for community assembly. Moreover, this end state can be achieved by a (potentially non-unique) sequence of single species invasions.

### A New Approach to combine modern coexistence theory with permanence theory

In the previous section, we discussed how we can use invasion growth rates of all possible sub-communities, via the invasion graph, to characterize all possible paths of community assembly, identify end states of community assembly, and improve our assessment of coexistence and invasion resistance for these end states. However, the main achievement of MCT is not assessing *whether* species can coexist, but rather unifying our understanding of *why* species coexist.

In principal, we could directly apply the methods from MCT to the entire invasion scheme, however, it is less clear what additional insight we might gain from this (but see Box 2), let alone how we would integrate across all of this information to make statements about the relative importance of different coexistence mechanisms at the scale of the community. Specifically, the potential number of invasion growth rates grows super-exponentially (up to 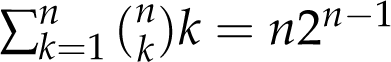 invasion growth rates), e.g. for a 10 species community we could have up to 5, 120 invasion growth rates. Rather, we suggest to focus on a specific subset of all these invasion growth rates, the ones which potentially hold the most information about the coexistence of the community and are best linked to current practices of MCT.

Suppose, for the moment, that all *n* species are able to coexist. Then, we propose to focus on the invasion growth rates of all potentially persistent communities with *n* − 1 species. These invasion growth rates focus one’s attention on the most proximate causes of coexistence. Indeed, for the full community to assemble via single species invasions, the final step of assembly would involve the invasion of the missing species from a community with *n* − 1 species (see Appendix Appendix E and Box 2 for alternative choices).

But what if not all *n* species coexist? Any collection of *n* species with an acyclic invasion graph has at least one community *S* corresponding to an end state of community assembly. To understand the coexistence of the species in *S*, we apply approach above replacing “*n*” with the number of species *k < n* in *S*. Furthermore, to understand why this community *S* is invasion resistant, we propose to apply the approach of MCT to the invasion growth rates of missing species. As these invasion growth rates *r_i_*(*S*) will be negative, the methodology of MCT is used to understand what contributes most to these negative values.

### Application to empirical data

We now focus on examples of the modern coexistence literature where our approach would give us additional insight that was not available with the naïve invasion criterion. We here assume that readers are familiar with the methods to compute niche and fitness differences or relative non-linearities and storage effect (Spaak *et al*., 2021b; Ellner *et al*., 2019). There are many different methods to compute niche and fitness differences (see Spaak *et al*. (2022b) for a review) and multiple methods to assess storage effect and relative non-linearity (Chesson, 2003; Barabás *et al*., 2018; Ellner *et al*., 2019). We chose the methods of Spaak *et al*. (2021b) as it does not rely on a pairwise comparison of species and Ellner *et al*. (2019) as it is easiest to apply to complex communities.

#### Fitness versus niche differences in competing plants

Spaak *et al*. (2021a) investigated how species richness affects niche and fitness differences in 33 empirical multispecies Lotka-Volterra communities. For many of these communities they were not able to compute niche and fitness differences, as the *n* − 1 communities did not coexist.

We focus on a six species community of competing plants (Geijzendorffer *et al*., 2011) and compute their niche and fitness differences (see appendix Appendix C). The invasion graph for this community is acyclic (Fig. 3A) and has a unique end state consisting of species {1, 2, 4, 5, 6}. However, for this end state not all *n* − 1 = 4 subcommunities exist, therefore Spaak *et al*. (2021a) did not compute niche and fitness differences for this end state (Fig. 3C). With our new approach, we focus solely on the positive invasion growth rates of species 1, 2 and 5, as these species define the only *n* − 1 subcommunities of the {1, 2, 4, 5, 6} community. We compute the niche and fitness differences of the competing species 1, 2 and 5, in presence of the other species - {1, 2, 4, 6} for species 5, {1, 4, 5, 6} for species 2 and {2, 4, 5, 6} for species 1 (Fig. 3D). For each these species, coexistence is mainly possible due to large niche differences, including strong facilitation for species 5. In contrast, the fitness difference component of species 3’s invasion growth rates into the resident community exceeds the niche difference component (Fig. 3D). Consequently, species 3 is excluded. We therefore understand that the coexisting species can coexist due to strong facilitation and unique niches, conversely, the excluded species is excluded due to too low niche differentiation, not due to low competitive ability.

#### Fluctuating coexistence mechanisms for floral yeast communities

Letten *et al*. (2018) studied coexistence for four yeast species competing for amino acids in flowers with temporally fluctuating sugar concentrations. They compute relative nonlinearity and storage effect for all six pair-wise combinations of species. We here repeat their analysis, but focus on the entire community at once. Consistent with Letten *et al*. (2018), the invasion graph of this community (Fig. 4A) has only one end state consisting of species 1 and 2. Coexistence of 1 and 2 reduces to the usual mutual invasibility criteria i.e. both *n* − 1 subcommunities of the *n* = 2 community {1, 2} exist. Hence, the understanding of why species 1 and 2 coexist remains the same as Letten *et al*. (2018)’s analysis of the {1, 2}.

Our approach potentially changes how we interpret the exclusion of species 3 and 4. To understand why species 3 and 4 are excluded we decompose the invasion growth rates *r_i_* into fluctuation independent effects (Δ_*i*_^0^) and fluctuation dependent effects. Furthermore, we decomposed the effects of fluctuating sugar concentration and fluctuating resource concentrations into relative non-linearities (Δ_*i*_*^S^* and Δ_*i*_*^R^*) and *i i* storage effect (Δ_*i*_(*^R#S^*) and Δ_*i*_*^(RS)^*) using the method of Ellner *et al*. (2019). However, depending into which resident community *S* they invade our understanding of their exclusion may change. Letten *et al*. (2018) focused on pair-wise coexistence and therefore analyzed the invasion growth rates of species 3 and 4 into the resident communities consisting of only species 1 (blue bars) or only species 2 (green bars) i.e. the computed *r_i_*(*S*) with *i* = 3, 4 and *S* = {1} or *S* = {2}. For our approach, we compute invasion growth rates *r*_3_(*S*), *r*_4_(*S*) into the combined community (*S* = {1, 2}) of species 1 and 2 coexisting (orange bars). For species 3 and 4 the positive storage effect Δ_*i*_^(*R#S)*^ and the positive relative non-linearity for the resource concentration (Δ_*i*_*^R^*) are negated by the negative relative non-linearity for sugar concentration (Δ_*i*_*^S^*) leading to an invasion growth rate *r_i_* which is approximately equal to the fluctuation independent growth rate (Δ_*i*_^0^). Additionally, we see that the coexistence mechanisms for the invasion into the sub-community {1, 2} are approximately the mean of the coexistence mechanisms for the invasion into the sub-community {1} and the sub-community {2}.

## Discussion

Modern coexistence theory (MCT) offers reliable tools to understand why species do or do not coexist in two-species communities or communities with diffuse inter-actions, but so far lacked tools to understand what limits diversity in species rich communities. As such, MCT allowed understanding of only one side of the biodiversity coin in species rich communities. Our proposed changes hold the potential to extend our understanding in two ways. First, we separate the process of assessing *whether* a community coexists from understanding *why* this community coexists. This allows us to focus on only the invasion growth rates of the existing *n* − 1 subcommunities without having to worry about the non-existing *n* − 1 subcommunities, the most frequent issue in applications of MCT to multi-species communities (Saavedra *et al*., 2017; Spaak *et al*., 2021a). Second, for a non-coexisting species *j* we apply the methods of MCT to the invasion growth rate of species *j* into this coexisting subcommunity.

Our new approach has different implications for three different community types; 1. Our new approach does not change any analysis for a community where all *n* species coexist and all *n* − 1 communities exist (e.g. Fig 2 A), which might be the majority of the existing empirical data used in MCT (Buche *et al*., 2022). 2. Our new approach allows the analysis for a community where all *n* species coexist, but not all *n* − 1 communities exist (e.g. Fig. 2 B), which were previously left unanalyzed (Spaak *et al*., 2021a). 3. Potentially most importantly, for a community where only *k* of the *n* species coexist we can no analyze both why these *k* species coexist using MCT but we can also analyze why the other *n* − *k* species are excluded and understand not only coexistence, but also non-coexistence (e.g. Fig. 2C, E, 3A and 4A). The understanding of the coexistence of the *k* species sub-community depends solely on the *k* species itself. The understanding of the non-coexistence of any other species *i* depends solely on the *k* species and the species *i* itself, i.e. the excluded species do not affect our understanding of (non-)coexistence of any of the other species.

MCT has helped us to understand how different drivers promote coexistence, such as phylogeny (Godoy & Levine, 2014; Narwani *et al*., 2013), functional traits (Kraft *et al*., 2015b; Gallego *et al*., 2019) or environmental conditions (Bimler *et al*., 2018). However, because of the limitations of MCT this work is almost exclusively done on two-species communities or communities with diffuse interactions, in part because MCT has been developed to understand how species from one guild can coexist. Consequentially, we know little about how multi-species communities coexist (Spaak *et al*., 2021a), especially from different guilds, and even less about why not more species coexist. For example, Godoy *et al*. (2014) analyzed the co-occurrence of 18 annual plant species and found that at most three of them can stably coexist according to MCT (Godoy *et al*., 2017). With our methods, one can analyze which of these communities are end-states and why the other species can not invade.

A better understanding of non-coexistence is direly needed, as often models fitted to empirical data predict competitive exclusion despite co-occurring species (Buche *et al*., 2022; Germain *et al*., 2016; Kraft *et al*., 2015b; Godoy & Levine, 2014). This mismatch between empirical observation may stem from difficulties of assessing the actual underlying species interactions (Adler *et al*., 2018a), from uncertainties (Bowler *et al*., 2022) or from a failure to consider the correct spatial dimension (Ricklefs, 2008; Hart *et al*., 2017). However, it is also conceivable that non-coexistence is more than solely the absence of coexistence. For example, alternative stable states are driven by positive frequency dependence, while coexistence is driven by negative frequency dependence (Ke & Letten, 2018; Schreiber *et al*., 2019; Mordecai, 2011) including the negative storage effect (Chesson, 1982, 1994; Schreiber, 2021, 2022). More generally, in a meta-analysis of niche and fitness differences across different ecological communities Buche *et al*. (2022) found that niche differences of coexisting species-pairs differ qualitatively from niche differences of competitively excluded species, independent of whether these were driven by positive or negative frequency dependence. However, most of these results stem from simple two-species communities and give little general insight to the multi-species context. Our newly proposed method would allow for a deeper investigation of why species do *not* coexist.

### Open problems

While our newly proposed method provides a mathematically rigorous approach to understanding coexistence, its application to species rich communities provides computational challenges. For example, to generate the invasion scheme, one has to identify the equilibria associated with all possible subcommunities i.e. all feasible equilibria (Saavedra *et al*., 2017). For the brute force method that we use, this requires solving the system for every possible subcommunity, a number which grows exponentially with the total number *n* of species. This is especially prohibitive for com-plex models, e.g. spatially explicit models. Importantly, while our method reduces the complexity of determining *n* species coexistence (in the sense of permanence) to finding the equilibria of all subcommunities, this is only possible if we have a simple way to find all of these equilibria.

There are two ways that one might approach this computational complexity, corresponding to a bottom-up community assembly and a top-down assembly (Serván & Allesina, 2021). In the bottom-up approach one may possibly devise algorithms that construct the invasion graph iteratively, rather than by brute force, by following the assembly process from less to more species rich communities. In the top-down assembly, advocated by Ellner *et al*. (2019), one may numerically simulate the model with all but the focal species *i* and verify that species *i* can re-invade this commu-nity (called −*i* community, Appendix A). However, in this approach one needs to be careful to identify all such communities and, consequently perform numerical simulations for multiple initial conditions. Additionally, in this approach we do not verify that the invasion graph is acyclic or that all communities with more species missing can indeed be invaded. None the less, under the assumption of being acyclic, the computation complexity only increases linearly with species richness. Hence, identifying food web topologies that ensure acyclic invasion graphs is an important open problem. For example, tritrophic food webs consisting of specialist predators and top-predators likely have acyclic invasion graphs (Wolkowicz, 1989).

We do not fully understand how to deal with invasion graphs containing cycles. We conjecture that coexistence occurs if there are appropriate weightings for each cycle which make the weighted average of the invasion growth rates positive along the cycle – the Hofbauer criterion (see Appendix B). However, it is unclear how the weightings from the Hofbauer criterion (see Appendix B) relate to the current metrics of MCT and whether we need a more generalized theory. Furthermore, unlike the acyclic case, the criterion in appendix Appendix B likely is sufficient for coexistence, but certainly not necessary. This discrepancy stems from a long-standing mathematical challenge of characterizing whether cycles, known as heteroclinic cycles in the dynamical systems literature, correspond to attractors leading to extinction or repellers aiding coexistence (Hofbauer, 1994; Brannath, 1994; Krupa, 1997). For a special class of “simple” cycles, Hofbauer (1994) has shown the sufficient conditions for coexistence are also necessary. The challenge of how to identify necessary criteria for coexistence beyond these simple cycles remains.

Our method ensures permanence, a very strong form of coexistence, ensuring that no species will go extinct even after a large perturbation. However, other definitions of coexistence, notably the existence of a positive attractor, are equally important (Schreiber, 2006). This is especially the case when species exhibit positive frequency dependence (Schreiber *et al*., 2017) or Allee effects (Stephens & Sutherland, 1999). When these mechanisms are operating, communities can exhibit alternative stable states supporting all species (coexistence) and subsets of species (non-coexistence). In these cases, invasion growth rates only help for identifying the latter alternative stable states, not the former. The theory of structural stability of feasible equilibria provides some tools for identifying the potential existence of coexistence at a stable equilibria (Saavedra *et al*., 2017). Coupling this theory with bifurcation theory (Guckenheimer & Holmes, 2013) provides a general approach to identifying non-equilibrium, positive attractors. However, equilibria-based bifurcations are not the only source of attractors of coexistence. Moreover, it’s unclear how to integrate these bifurcation methods into MCT (see, however, Barabás *et al*. (2012, 2014)).

## Conclusions

We have proposed a more general method to assess coexistence of a multi-species community. This method provides an essential link between MCT and community assembly. For many communities analyzed so far, where all species coexist and naïve invasion growth rate criterion is applicable, nothing has changed. However, with the method we can now apply the tools of MCT to species rich communities (Chesson, 2018; Spaak *et al*., 2021a) including multi-trophic communities (Spaak *et al*., 2021b; Shoemaker *et al*., 2020; Godoy *et al*., 2018) where the naïve criterion cannot be applied. This method also identifies the end states of community assembly when all species can not coexist and helps explain why end states are invasion resistant. Looking forward, our method may provide insights into how the balance of coexistence mechanisms shift during community assembly.

## Boxes

### Box 1

How to identify assembly end states

1. **Compute the invasion scheme:** For each possible sub-community *S* ⊂ {1, 2, …, *n*} numerically compute the long-term dynamics of the subcommunity i.e. an equilibrium (*N*^^^, *A*^^^) or an approximation of the stationary density *p_S_*(*dN*, *dA*) (see Appendix Appendix D) supporting these species (i.e. *N*^^^*_i_ >* 0 or ∫ *N_i_ p_S_*(*dN*, *dA*) *>* 0 if and only if *i* is in *S*). There are 2*^n^* – 1 possible sub-communities. For each sub-community compute the invasion growth rates *r_i_*(*S*) of the absent species.
2. **Generate the invasion graph:** The subcommunities from the invasion scheme are the vertices of the invasion graph. For each pair of subcommunities *T* and *S* where *S* ≠ *T*, add a directed edge from *S* to *T* if

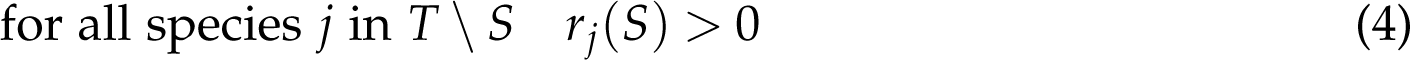

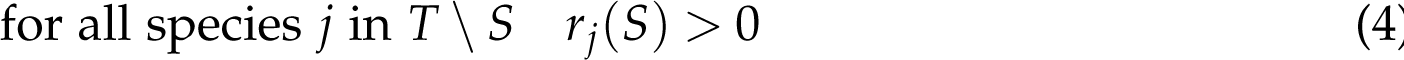 Intuitively, this implies that community *T* could be reached from community *S* by invasion.
3. **Check for cycles**: Determine whether the invasion graph is acyclic or not. This can be done, for example, using the is dag command from the igraph package in R (Csardi & Nepusz, 2006). If the graph is acyclic, continue to step 4, else use methods discussed in Appendix B.
4. **Find an end-state:** Find an *S* ⊂ {1, 2, …, *n*} such that a) *S* is uninvadable i.e. *r_i_*(*S*) *<* 0 for all species *i* not in *S*, and b) for every subcommunity *T* of *S*, there is at least one species in *S* that can invade *T* i.e. *r_i_*(*T*) *>* 0. Condition b) ensures that the subcommunity *S* is permanent.
5. **Analyze invasion growth rates ”***n* − 1**” subcommunities of** *S*: Coexistence of the end state *S* hinges on the invasion growth rates from the sub-communities *T* of *S* with exactly one less species than *S*. We therefore apply the methods of MCT to these decompose and partition these invasion growth rates. Importantly, we potentially analyze the persistence of only a sub-set of the maximal community.
6. **Analyze the invasion growth rates of excluded species (if any) of the assembly end state** *S*: If the end state *S* is a proper sub-community (i.e. has less than *n* species), then analyze the negative invasion growth rates of the competitively excluded species with the methods of MCT.

We provide automated code for step 1 for Lotka-Volterra community models, steps 2-4 for any model type, and steps 5-6 for niche and fitness differences for Lotka-Volterra community models (Appendix C).

### Box 2

Alternative uses of the invasion graph

In the main text we focused on applying the methods of MCT to the *n* − 1 communities, i.e. the last steps of the community assembly, for two reasons. First, after the loss of any species, reassembly of the community depends on the invasion growth rate into one of these *n* − 1 communities. Second, the resulting applications most resemble the current use of MCT. However, the invasion graph contains much more information. We offer three alternative applications of MCT to the invasion graph.

**What hampers the formation of missing** *n* − 1 **communities?** After identifying an end state without all of its *n* − 1 communities, we may ask why a certain species *i* does not have its corresponding *n* − 1 community. Equivalently, why doesn’t this *n* − 1 subcommunity coexist? As an example we take the community in Figure 3C. Here, species 4 does not have an *n* − 1 community, as the other species can not coexist. The other species can not coexist because species 5 has a negative invasion growth rate into the community *S* = {1, 2, 6}, so the community *S* is an end-state with respect to the species (1, 2, 5, 6). However, species 4 can invade *S* leading to the community *T* = {1, 2, 4, 6}. Importantly, species 5 has a positive invasion growth rate into this new community *T*. Here, the invasion of species 4 into *S* greatly increases the niche differences of species 5, but only slightly increases its fitness differences, leading to an overall positive invasion growth rate of species 5. Hence, species 4 indirectly facilitates species 5 and, consequently, the {1, 2, 5, 6} can not form in the absence of species 4. In general, we conjecture that the *n* − 1 associated with a species *i* only fails to form if species *i* is indirectly facilitating one of the other species in a similar way i.e. when species *i* invades the community *S* formed in its removal, it leads to a new community *T* that one of the other missing species in *S* can invade.
**How do the metrics of MCT change along community assembly?** Suppose we have a community of *n* species where all sub-communities exist (e.g. Fig. 2A with *n* = 3) and are interested in the assembly path ᴓ → {1} → {1, 2} → · · · → {1, 2, …, *n* − 1}. In this case we might ask how the invasion growth of the species *n* changes along this assembly path and how the different metrics of MCT are affected by increasing species richness. More generally, we might ask whether certain mechanisms are stronger or more important at the beginning of community assembly versus the end of community assembly. This may shed new light on the question of why community assembly eventually stops and what limits diversity. For example, in Figure 3, niche and fitness differences seem to exhibit greater variation at the start of the assembly process (Fig.3B) than at the end of the assembly process (Fig.3D); this seems to be true throughout the assembly process (Figure A3).
**Which mechanisms are responsible for the creation of assembly loops?** We mostly excluded invasion graphs containing cycles as the invasion graph may not capture all transitions in the ecological dynamics, assessing permanence becomes more challenging, and it unclear how best to apply the methods of MCT to them. However, we could apply the methods of MCT along a cycle to understand what drives the alternating positive and negative invasion growth rates along the cycle and, thereby, potentially help us understand coexistence in invasion graphs with cycles.

**Figure 3:**
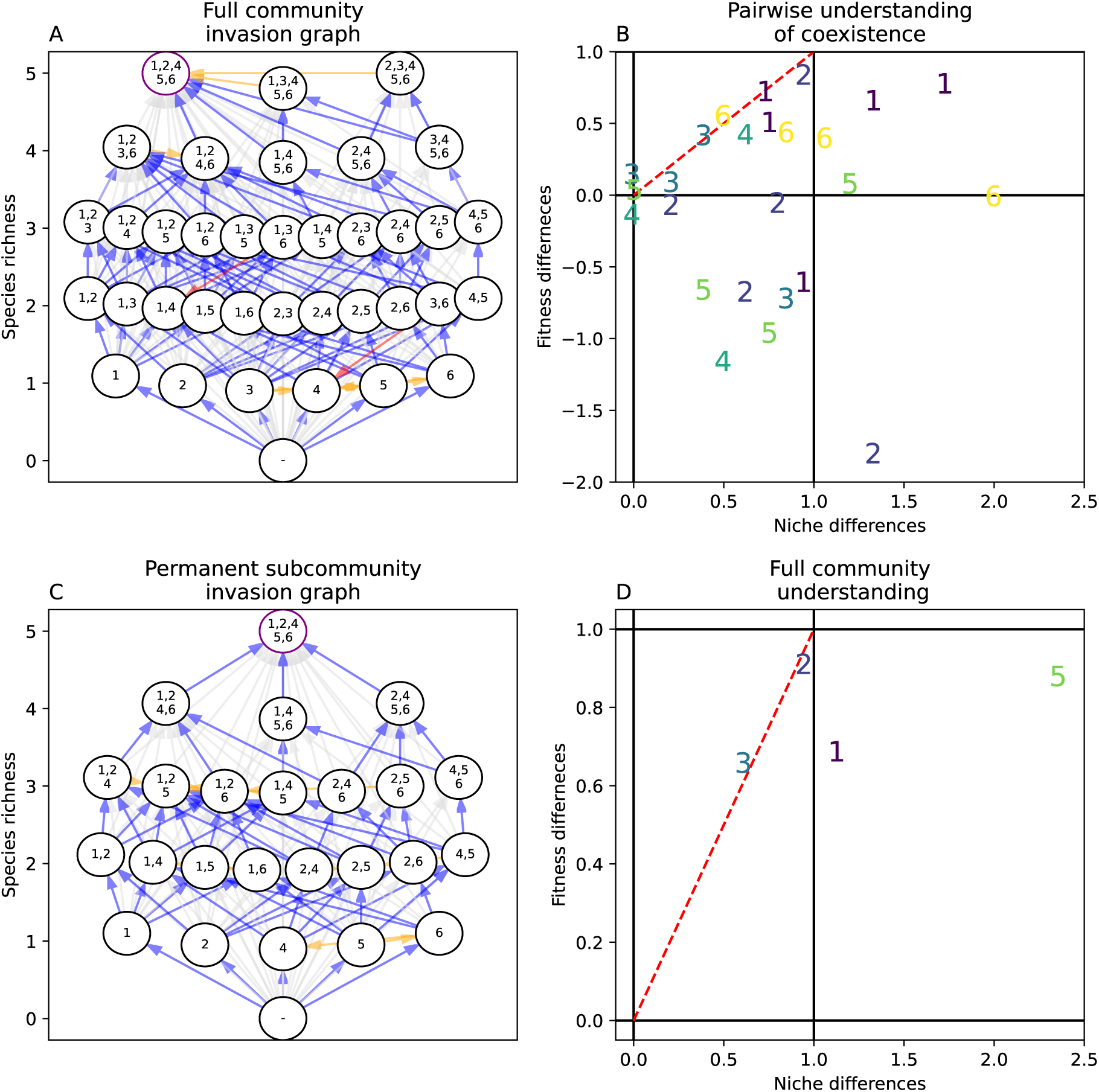
We applied the new method to a dataset from Spaak *et al*. (2021a). A: The acyclic invasion graph of the non-permanent six species community where 1, 2, 4, 5, 6 is a permanent and invasion-resistant sub-community. B: Assessing pairwise coexistence reveals that species 3 is excluded by species 4 and species 5, species 5 is excluded by species 6, and species 6 is excluded by species 4. This pair-wise analysis does not allow us to draw any conclusions of why species 3 is excluded from the full community. C: For the invasion graph in (A) restricted to 1, 2, 4, 5, 6. Only the removal of species 1, 2 and 5 define *n* 1 communities. D: For species 1,2 and 5 niche differences are strong enough to overcome fitness differences.Coexistence of the species 1, 2 and 5 is mainly possible due to large niche differences, including strong facilitation for species 5. Niche differences can not overcome fitness difference of the competitively excluded species 3. The other species (4 and 6) are not explicitly included into the coexistence analysis, as they do not form an *n* 1 community. The analysis of the entire community, compared to the pair-wise analysis shown in panel B, gives a more complete understanding of why species 3 is excluded: it has the lowest fitness differences but not sufficiently 1s5trong niche differences.

## Appendix A Mathematical Assumptions, −*i* communities, and Extensions

In this Appendix, we describe the mathematical assumptions introduced in Hofbauer & Schreiber (2022) and discuss how these assumptions can be relaxed using techniques introduced in Schreiber (2000). As stated in the main text, we focus on continuous-time models of *n* interacting species with non-negative densities *N* = (*N*_1_, *N*_2_, …, *N_n_*). To allow for population structure (e.g. discrete habitat patches, stages, genotypes), temporal forcing (e.g. periodic or chaotic environmental fluctuations), and environmental feedbacks (e.g. plant-soil feedbacks), we allow for a finite number *m* of auxiliary variables *A* = (*A*_1_, *A*_2_, …, *A_m_*) which can be positive or negative. Many models often used in MCT do not have any auxiliary variables, Lotka-Volterra models and general Lotka-Volterra community models as well as many annual plant models (Levine & HilleRisLambers, 2009; Godoy & Levine, 2014). Conversely, resource competition models have the resource densities as auxiliary variables (Letten *et al*., 2017) and the lottery model, as well as many similar models have the environment as an auxiliary variable (Chesson, 1994). The use of auxiliary variables is discussed in (Patel & Schreiber, 2018a; Benaïm & Schreiber, 2019).

The per-capita growth rate *f_i_*(*N̂*, *Â*) of species *i* depends on both the species’ densities and auxiliary variables. The rate of change *g_j_*(*N̂*, *Â*) of the *j*-th auxiliary variable also depends on both densities and auxiliary variables.

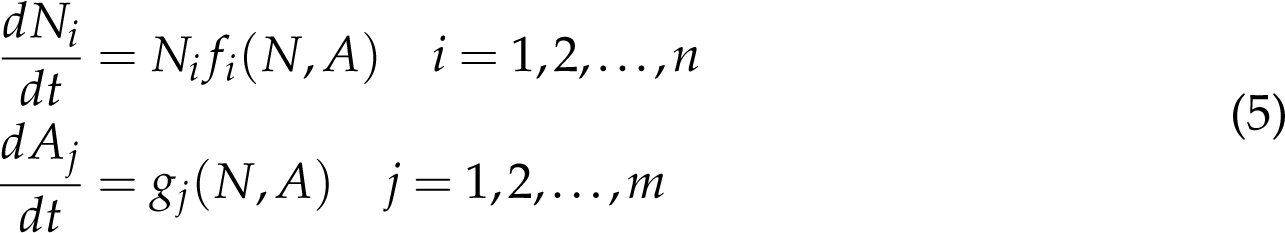

### Main assumptions of the theory

The results from Hofbauer & Schreiber (2022) are based on four main assumptions. We lay out these assumptions mathematically, and then provide an intuitive interpretation of these mathematical statements. These interpretations are meant to give an intuitive understanding of the theorem.

1. the functions *f_i_*(*N̂*, *Â*) are *g_j_*(*N̂*, *Â*) are continuously differentiable. Namely, the rates of change (*f_i_* and *g_i_*) and their derivatives vary continuously with the species densities and auxiliary variables. Importantly, this assumption ensures that there exist a unique solution (*N*(*t*), *A*(*t*)) to (5) for any initial condition (*N*(0), *A*(0)). Intuitively, this means that *f_i_*and *g_j_*are “nice” functions and that we can indeed talk about densities over time. Most reasonable, biological function satisfies this mathe-matical assumption.
2. the system (5) is dissipative: there is some positive constant *K >* 0 such that every solution (*N*(*t*), *A*(*t*)) satisfies *N_i_*(*t*) ≤ *K* and |*A_j_*(*t*)| ≤ *K* for all *i*, *j* and *t* ≥ 0 sufficiently large. Intuitively, this implies that the species densities and the auxiliary variables are eventually bounded. Counterexamples do exist, but they are typically not biologically realistic, e.g. facilitation leading to an explosion of densities or unlimited evolution allowing infinitely large species.

The final two assumptions concern stationary distributions *p*(*N̂*, *Â*)*dNdA* that are ergodic: a stationary distribution that can not be written as a non-trivial convex combination of two other stationary distributions. Ergodicity of a stationary distribution implies there is a unique set of species, denoted *S*(*p*) ⊂ {1, 2, …, *n*} (“the support of *p*”), such that _A_ *p*(*N̂*, *Â*)*dNdA* = 1 where A = {(*N̂*, *Â*) : *N_i_ >* 0 iff *i* ∈ *S*(*p*)}. In words, *S*(*p*) is the set of species whose densities are positive at this distribution. Using an argument introduced in (Schreiber, 2000), Lemma 1 of (Hofbauer & Schreiber, 2022) implies that the average per-capita growth rate of species supported an ergodic stationary distribution equal zero i.e. *r_i_*(*p*) = *f_i_*(*N̂*, *Â*)*p*(*N̂*, *Â*)*dNdA* = 0 for all *i* ∈ *S*(*p*). Intuitively, if a species densities are bounded from above and away from zero, then in the long term its average per-capita growth rate is zero.

1. the invasion growth rates of all the missing species *j* ∈/ *S*(*p*) are non-zero i.e. *r_i_*(*p*) ≠ 0. The proof of (Schreiber, 2000, Theorem 4.1) suggests that this assumption is often met. Two neutral species would violate this assumption, but this is a nongeneric, yet theoretically instructive, situation. Hening *et al*. (2022) proved that this assumption is generic for stochastic differential equation models.
2. The most important assumption is that the signs of the invasion growth rates are consistent among all communities represented by an ergodic stationary distribution. More precisely, if *p*(*N̂*, *Â*)*dNdA* and *q*(*N̂*, *Â*)*dNdA* are two ergodic stationary distributions supporting the same species (i.e. *S*(*p*) = *S*(*q*)) then

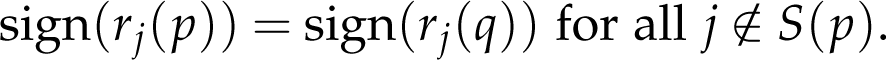
3. Intuitively, this assumption implies that if we know which species are present, then we know which absent species can invade. This assumption is automatically true if for each combination of species there is only one stationary distribution.

Overall, assumptions 1-3 are technical mathematical assumptions that likely are meet for most models. However, assumption 4 is not purely a technicality and can be violated by reasonable biological systems. Importantly, all four assumptions are met for general Lotka-Volterra models. An important class of models for which that these assumptions are naturally met are general Lotka-Volterra models where the per-capita growth rates are linear functions of the species densities i.e. *f_i_*(*N*) = ∑*_j_ a_ij_n_j_* + *b_i_*. For dissipative Lotka-Volterra models (see, e.g., Hofbauer & Sigmund (1998, Theorems 15.2.1,15.2.4) for sufficient conditions), the remaining three assumptions are generically met i.e. for arbitrarily small perturbations of *A* and *b*, one can ensure that the three assumptions are met. This follows from the time averaging property of Lotka-Volterra systems (Hofbauer & Sigmund, 1998, Theorem 5.2.3) and that generically the per-capita growth rates at equilibria satisfy the third assumption.

For non-Lotka Volterra systems, however, it is possible for the per-capita growth rates of a missing species to have opposite signs at different ergodic stationary distributions. For example, this failure arises in models of two predator species competing for a single prey species (McGehee & Armstrong, 1977). If one predator has a type II functional response, then the predator-prey subsystem may simultaneously have an unstable equilibrium (defining one ergodic stationary distribution) and a stable limit cycle (defining a nother e rgodic s tationary d istribution). McGehee & Armstrong (1977) showed that the invasion growth rates of the other predator species may be positive at the stable limit cycle but negative at the unstable equilibrium. A similar phenomena arises in models of two prey species sharing a common predator (Schreiber, 2004).

### What *S* → *T* means dynamically

A key result proved by Hofbauer & Schreiber (2022) highlights what aspects of the community dynamics are captured by the invasion graph. Roughly, this result states that if a solution (*N*(*t*), *A*(*t*)) converges in forward time to an equilibrium, periodic orbit, etc corresponding to community *T*, but in backward time converges to an equilibrium, periodic orbit, etc. corresponding to community *S*, then the invasion graph includes the transition *S* → *T*. More precisely, one can define the *ω*-limit set of the solution (*N*(*t*), *A*(*t*)) to be the set of points (*N̂*, *Â*) such that (*N*(*t_k_*), *A*(*t_k_*)) → (*N̂*, *Â*) as *k* → ∞ for some sequence of times *t*_1_, *t*_2_, … with *t_k_*→ ∞. The *α*-limit set of this solution is defined similarly except that *t_k_* → −∞. If for a solution (*N*(*t*), *A*(*t*)) its *ω*limit set is supported by the species in *T* and its *α*-limit set is supported by the species in *S*, then Hofbauer & Schreiber (2022) proved that the invasion graph includes the transition *S* → *T*.

### The coexistence criterion and −*i* communities

When the four assumptions are met, the invasion graph can be defined by only considering the signs of invasion growth rates associated with communities supported by at least one ergodic stationary distribution. For this reason, the invasion scheme is formally only defined by the signs of invasion growth rates, not their actual values. Hofbauer & Schreiber (2022, Theorem 1) proves that an acyclic invasion graph is permanent if and only there is a positive invasion growth rate for at least one missing species at every ergodic stationary distribution supporting a proper subset of species.

This result can be rephrased in terms of −*i* communities, a fundamental concept of coexistence theory (Chesson, 1994). −*i* communities are the sub-communities that form after species *i* is lost from the full community. In the context of invasion graphs, a −*i* community *S* is a community without species *i* and that resists invasion attempts from all other missing species i.e. *r_j_*(*S*) *<* 0 for *j* ≠ *i* and *j* ∈/ *S*. Under suitable technical assumptions (Hofbauer & Schreiber, 2022), for every −*i* community there are solutions (*N*(*t*), *A*(*t*)) of the community dynamics such that all species are initially present and in forward time the solution approaches an equilibrium (more generally, an ergodic stationary distribution) corresponding to *S*. Hofbauer & Schreiber (2022, Corollary 1) proved for acyclic invasion graphs, if *r_i_*(*S*) *>* 0 for all −*i* communities *S* and all species *i*, then the model is permanent i.e. all species *i* must be able to invade all −*i* communities for coexistence.

### Going beyond invasion graphs: Morse decompositions

To relax the assumptions of the invasion graphs, one needs to introduce a different type of graph that, in general, is not determined solely by the invasion growth rates (Schreiber, 2000; Roth *et al*., 2017; Patel & Schreiber, 2018a). Specifically, one needs the concept of a Morse decomposition for the dynamics on the extinction set i.e. (*N̂*, *Â*) where *N_i_*= 0 for at least one *i*. Roughly (see Schreiber (2000); Patel & Schreiber (2018a) for a precise definition), the Morse decomposition consists of a finite collection of compact, invariant sets *M*_1_, *M*_2_, … *M_k_* (e.g. equilibria, limit cycles, quasi-periodic motions, chaotic saddles, etc) that satisfy two properties.

1. all the long-term dynamics are contained in the union of these sets i.e. all *ω*limit sets for the extinction set lie in ∪*_i_ M_i_*. Intuitively, this implies that the community dynamics eventually approach one of the *M_i_*.
2. for any initial condition where in backward time the solution approaches *M_i_* and in forward time the solution approaches *M_j_*, *j* ≥ *i* with equality only if the initial condition was in *M_i_*.

Associated with this Morse decomposition is a directed graph where the vertices are the sets *M*_1_, …, *M_k_* and there is a directed edge from *M_i_* to *M_j_* (where *i* ≠ *j*) only if there is a solution which in backward time goes to *M_i_*and in forward time goes to *M_j_*. By the definition of the Morse decomposition, this graph is acyclic. A sufficient condition for coexistence, in the sense of permanence, is that for each component *M_i_* there is a species *j* such that their invasion growth rate is positive at every ergodic stationary distribution supported by *M_i_*. In the special case that the Morse components always lie in the interior of the extinction faces, this sufficient condition can be viewed as a generalization of the invasion graph approach of Hofbauer & Schreiber (2022).

## Appendix B Invasion Graphs with Cycles

Our work focuses on the case where the invasion graph contains no cycles. Cycles appear to be rare in the empirical data investigated by modern coexistence so far. Specifically, we only found one data set with cycles among many investigated communities (Chu & Adler, 2015; Spaak *et al*., 2021a; Adler *et al*., 2018b; Zepeda & Martorell, 2019; Godoy & Levine, 2014; Shoemaker *et al*., 2020; Letten *et al*., 2018), see also (Godoy *et al*., 2017). However, almost all these communities focused only on species in the basal trophic level, communities with multiple trophic levels might be very different (Schreiber & Rittenhouse, 2004; Spaak *et al*., 2022a; Law & Morton, 1996b; Song *et al*., 2021).

In invasion graphs with cycles, a simple way to assess permanence in applying the Hofbauer (1981) criterion to the invasion growth rates associated with all stationary distributions. Specifically, the Hofbauer criterion requires finding positive weights *w_i_* associated with each species such that

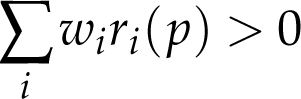

for every ergodic, stationary distribution *p*(*dN*, *dA*) supporting a strict subset *S*(*p*) ≠ {1, …, *n*} of species. Intuitively, ∑*_i_ w_i_r_i_* is a community-level invasion growth rate. If this community-level invasion growth rate is positive whenever one or more species is rare, then the community tends to recover and all the species coexist. However, the Hofbauer criterion is not a necessary condition for coexistence, only sufficient condition. That is, if such weightings *w_i_* exist such that ∑*_i_ w_i_r_i_*(*p*) *>* 0, then the community is permanent, however, the community can be permanent and no such weighting exists.

If no weight *w_i_* exist that work for all stationary *p*, then we conjecture that one can verify permanence by applying the Hofbauer criterion separately to stationary distributions associated with certain cycles in the invasion scheme. More precisely, we partition the vertices of the invasion graph into distinct subsets *V*, and the subcommunities *S* and *T* belong to the same subset if there is a cycle in the invasion graph which contains these two subcommunities. In other words, there is a sequence of invasion that go from *S* to *T*, and vice-versa. We then apply the Hofbauer cri-terion to the set of ergodic stationary distributions *p* associated with each of these subsets *V*. Namely, we must find weightings *w_i_^V^* such that ∑*_i_ w_*i*_^V^r_i_*(*p*) *>* 0 for all ergodic stationary distributions *p*(*dN*, *dA*) supporting communities *S*(*p*) lying in *V*. Intuitively, such a weighting *w_*i*_^V^* ensures that we will eventually exit the cycles in *V* and continue our path ”upwards” in the invasion graph. We conjecture that if such a weighting exists for all cycles, then we will eventually leave all these cycles and the entire community assembles. However, this condition would only be sufficient and not necessary for a permanent community.

These two applications of the Hofbauer criterion can at best assess the permanence of the community. To have a better understanding of why the community coexists one may then apply modern coexistence theory, with caution. If there exist *n* − 1 communities and none of the cycles contain these *n* − 1 communities, then we propose to apply modern coexistence theory to the *n* − 1 communities as discussed in Box 1. In essence, the cycles are only important during early phases of the community assembly, but they are not the bottlenecks of coexistence. However, it is also possible that none of the *n* − 1 communities exist (Fig. 2 D). How we can generalize the methods of modern coexistence theory to the Hofbauer criterion and cycles remains an open question.

## Appendix C Niche and fitness differences

We want to compute the niche and fitness differences for a Lotka-Volterra community with *n* species, given by 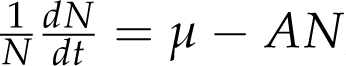, where *µ* is the vector of intrinsic growth rates, *A* is the species interaction matrix and *N* is the vector of species densities. We assume that the invasion graph is cycle free, that the community *S*, with *k* species, is permanent and that no species *i* not in *S* can invade into the community *S*. Finally, we denote the *n* − 1 communities of the community *S* as *T_j_*, i.e. *T_j_*consists of all *k* − 1 species in *S* except species *j*. We denote *J* as the set of *j* for which *T_j_* exists. The species are therefore separated into 3 groups: First, the species in *S* for which a *n* − 1 community exists, denoted *J*, second, the species in *S* for which a *n* − 1 community does not exist, denoted *S* − *J* and finally the species not in *S* denoted *E*, as they are excluded.

In the example given in figure 4 A this would imply *J* = {1, 2}, *T*_1_ = (2) and *T*_2_ = (1), *S* − *J* = {} and *E* = {3, 4} In the example given in figure 3 A this would imply *J* = (1, 2, 5), *T*_5_ = (1, 2, 4, 6), *T*_2_ = (1, 4, 5, 6) and *T*_1_ = (2, 4, 5, 6). *S* − *J* = {4, 6} as the subcommunities *T*_4_ and *T*_6_ do not exist and *E* = {3}. Note that the the communities (1, 2, 3, 6) and (3, 4, 5, 6) are not *n* − 1 communities of the community *S*, as they contain the species 3 which is not in *S*.

**Figure 4:**
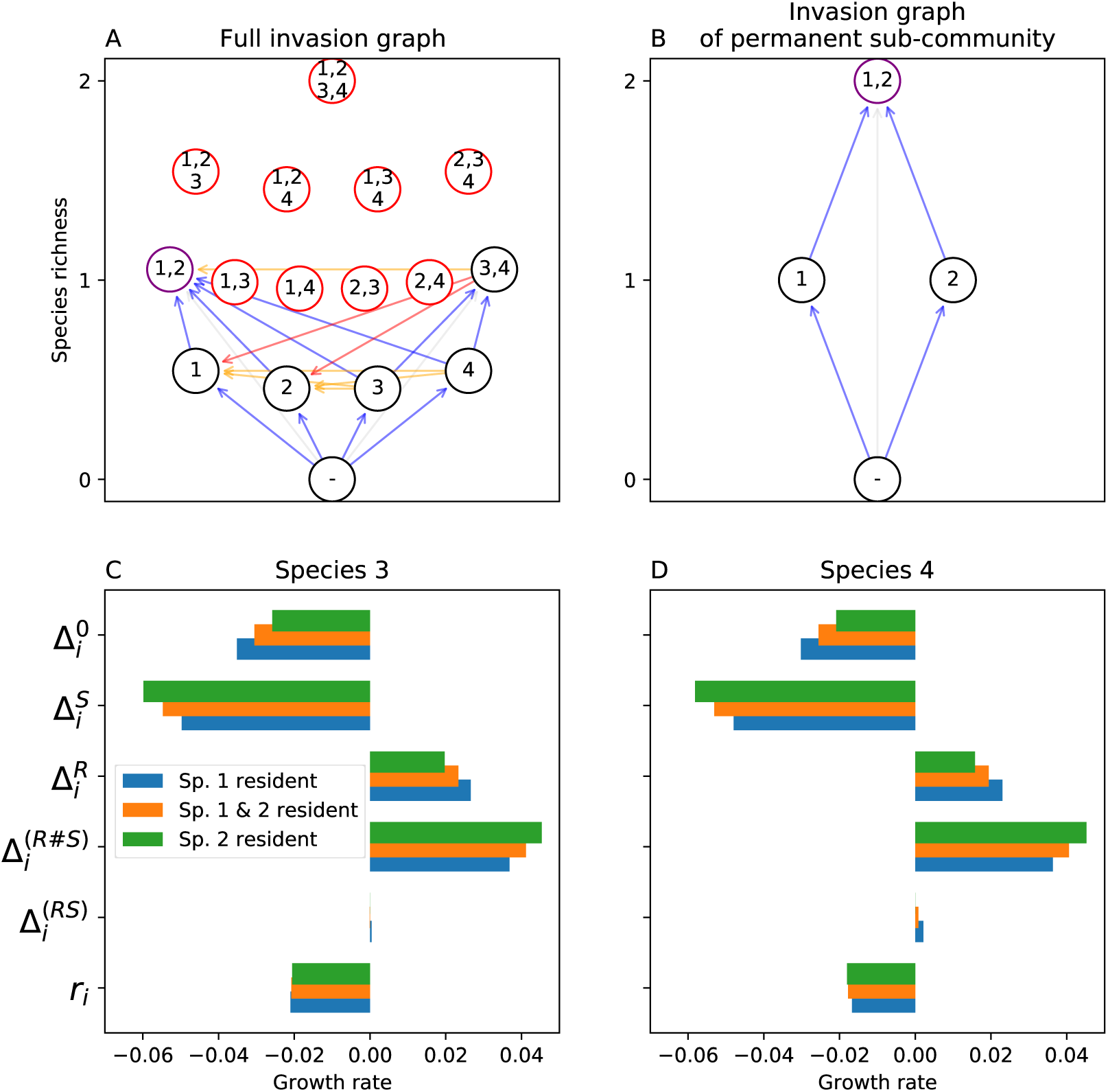
Letten *et al*. (2018) analyzed the coexistence of four competing yeast species which are not permanent (A). Species 1 and 2 can coexist and exclude species 3 and 4. Naïve invasion growth rate approach would explain their exclusion by computing their invasion growth rates into the subcommunities consisting of species 1 or species 2 only (green and blue bars). However, more informative is the decomposition of the invasion growth rates into the subcommunity consisting of species 1 and 2 (orange bars).

Given these groups we separate the intrinsic growth rate and the interaction ma-trix into these different groups, denoted *µ^X^* for the intrinsic growth rates of species in group *X* and *A^X^,^Y^* for the per-capita effects of species in group *Y* on species in group The dynamics of the community model is then given by

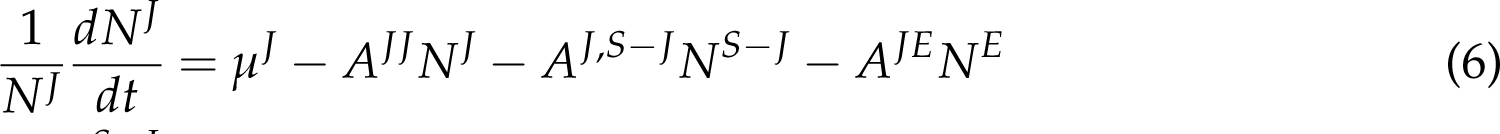

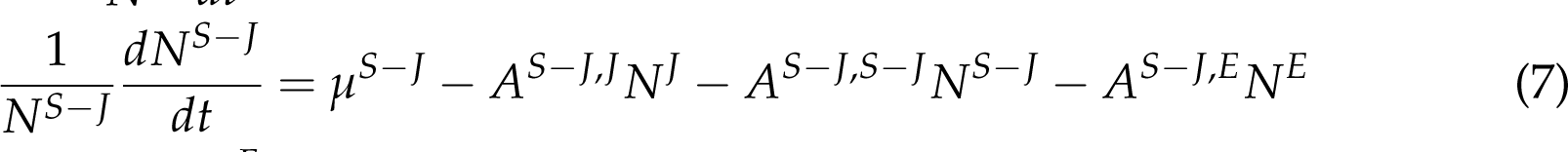

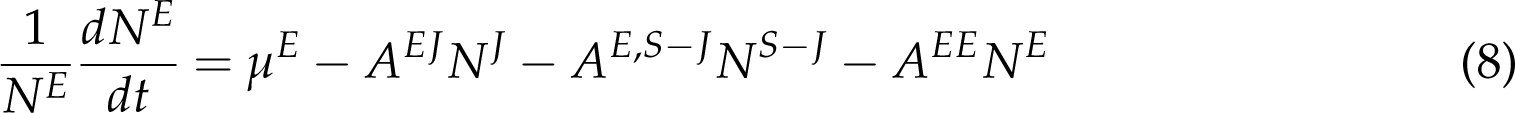

To compute niche and fitness differences we have to compute the intrinsic, invasion and no-niche growth rates. All of these growth rates are computed at real or hypothetical equilibria. When computing niche and fitness differences for the species *j* in *J* we have *N^E^* = 0, we therefore omit them from the model. The species *i* in *S* − *J* are treated similar to limiting factors, as they are not the focal species, as mentioned in the main text. We therefore solve eq. 7 for *N^S^*^−^*^J^* and get *N^S^*^−^*^J^* = *A^S^*^−^*^J^,^S^*^−^*^J^* ^−1^ *µ^S^*^−^*^J^* − *A^S^*^−^*^J^,^J^ N^J^*, which we can insert into eq. 6 to get

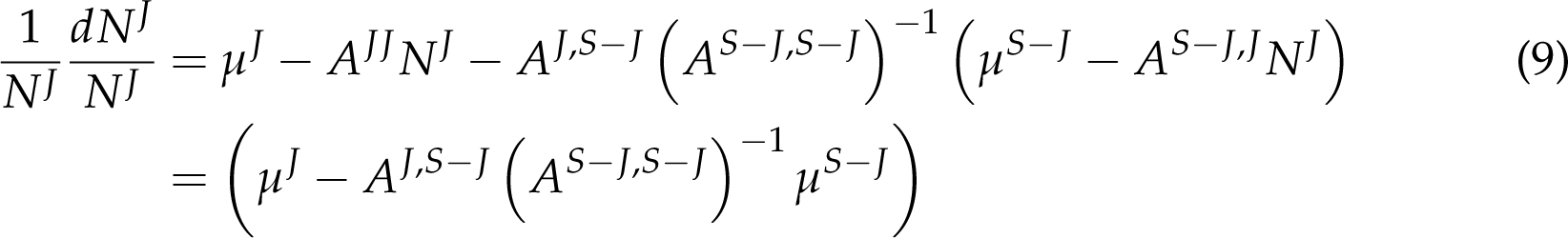

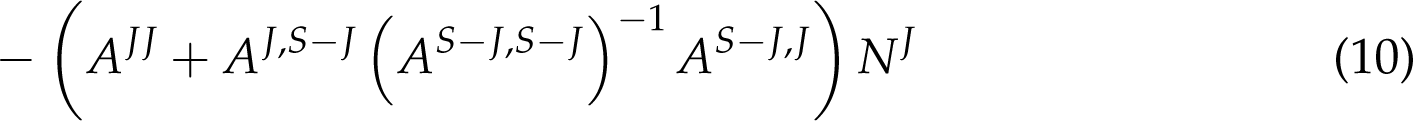

While appearing complicated, this is actually again a simple Lotka-Volterra commu-nity model with *_µ_*′ _=_ *_µ_J* _−_ *_A_J,S*−*J _A_S*−*J,S*−*J* ^−1^ *_µ_S*−*J* _and_ *_A_*′ _=_ *_A_JJ* _+_ *_A_J,S*−*J _A_S*−*J,S*−*J* ^−1^ *_A_S*−*J,J*. Importantly, for this new Lotka-Volterra system all species can coexist and all *n* − 1 communities exist, we can therefore compute niche and fitness differences as usual (Spaak *et al*., 2021a). This method is mathematically equivalent to the well-known time-scale separation (Chesson, 1990), however, it does not depend on the assumption of a time-scale separation. Rather, as the growth rates are only evaluated at equilibria of the system we can use the fact that 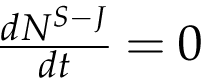 (Chesson & Kuang, 2008). To compute niche and fitness differences we have to compute the invasion growth rate *r_i_* = *µ_i_*^′^ − *A*^′^ *N^J^*^,−^*^i^*, where *N^J^*^,−^*^i^* is the equilibrium density of the resident community. Additionally, we have to compute the hypothetical invasion growth rate where all species interactions are absent, which is equivalent to the growth rate *µ_i_*^′^. Finally, we have to compute the no-niche growth rate where all niche differences of species *i* are removed. This is achieved by multiplying the actual inter-specific interaction strength with the two-species niche over-lap 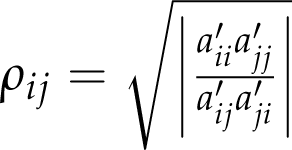, i.e. 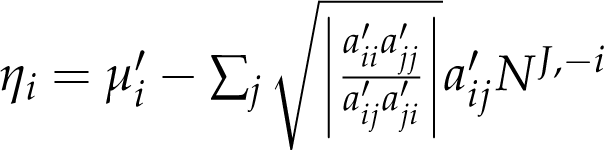. Note that the scaling of the inter-specific interaction coefficients results in no niche differences in the pair-wise community, but does not affect their fitness differences. Given these three growth rates we can then compute the niche and fitness differences as

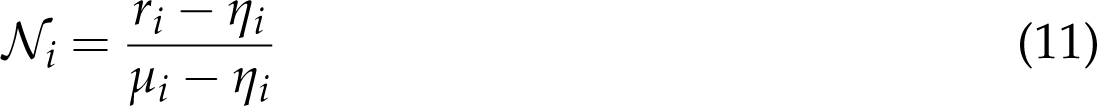

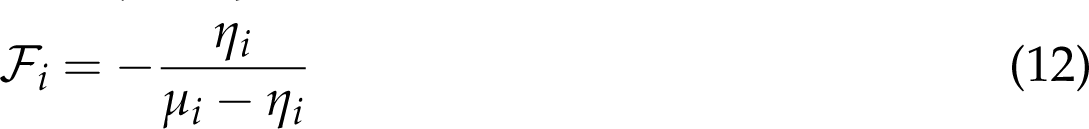

For the excluded species we can compute the intrinsic growth rates and the invasion growth rates as usual, however the invasion growth rates are defined into the resident community *S*, opposed as the usual invasion growth rates into an *n* − 1 community. The no-niche growth rate can be computed using the same scaling of the inter-specific interaction coefficients as discussed above (Spaak & De Laender, 2020).

## Appendix D Stationary distribution

Long-term ecological dynamics of a subcommunity *S* may exhibit various forms of non-equilibrium dynamics including periodic oscillations, quasi-periodic motions, chaos, and stochastic fluctuations. These non-equilibrium dynamics may be endogenously driven due species interactions, exogenously driven due to environmental forcing, or due to a combination of these effects. For example, Figure A1 illustrates a predator-prey subcommunity *S* (with densities *N*_1_, *N*_2_) experiencing seasonal forcing of the prey’s carrying capacity (determined by auxiliary variable *A*). The long-term statistical behavior of these non-equilibrium subcommunity dynamics are often characterized by a stationary distribution (*p_S_*(*N̂*, *Â*)*dNdA*) describing the fraction of time, in the long-term, that the system is in any given configuration. These stationary distributions, as we discuss more explicitly below, can be approximated by the (multivariate) histogram of a sufficiently long time series of the species densities and the auxiliary variables.

The invasion growth rate of species *i* is the average per-capita growth rate with respect to this stationary distribution *p_S_*(*dN*, *dA*):

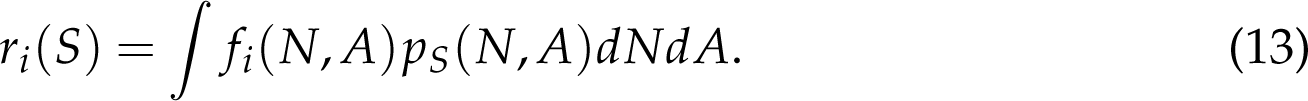

when the stationary distribution cannot be expressed as an average of other stationary distributions (see ergodicity in appendix Appendix A), Birkhoff’s ergodic theorem implies that

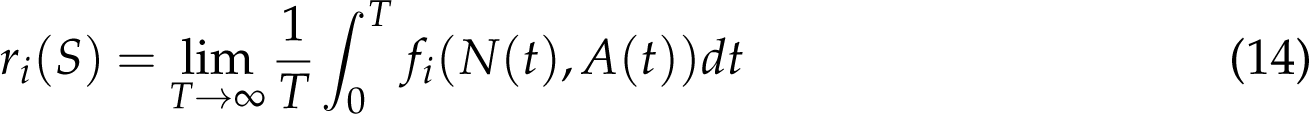

with respect to a randomly chosen initial condition (*N*(0), *A*(0)) from the stationary distribution. Hence, one can estimate the invasion growth rates *r_i_*(*S*) by (i) simulating the subcommunity *S* dynamics for a sufficiently long time *T* ≫ 1, (ii) computing the average per-capita growth rates 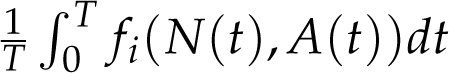 for each of the species *i* not in *S*, and (iii) setting *r_i_*(*S*) = 0 for all the species *i* in *S*.

**Figure A1:**
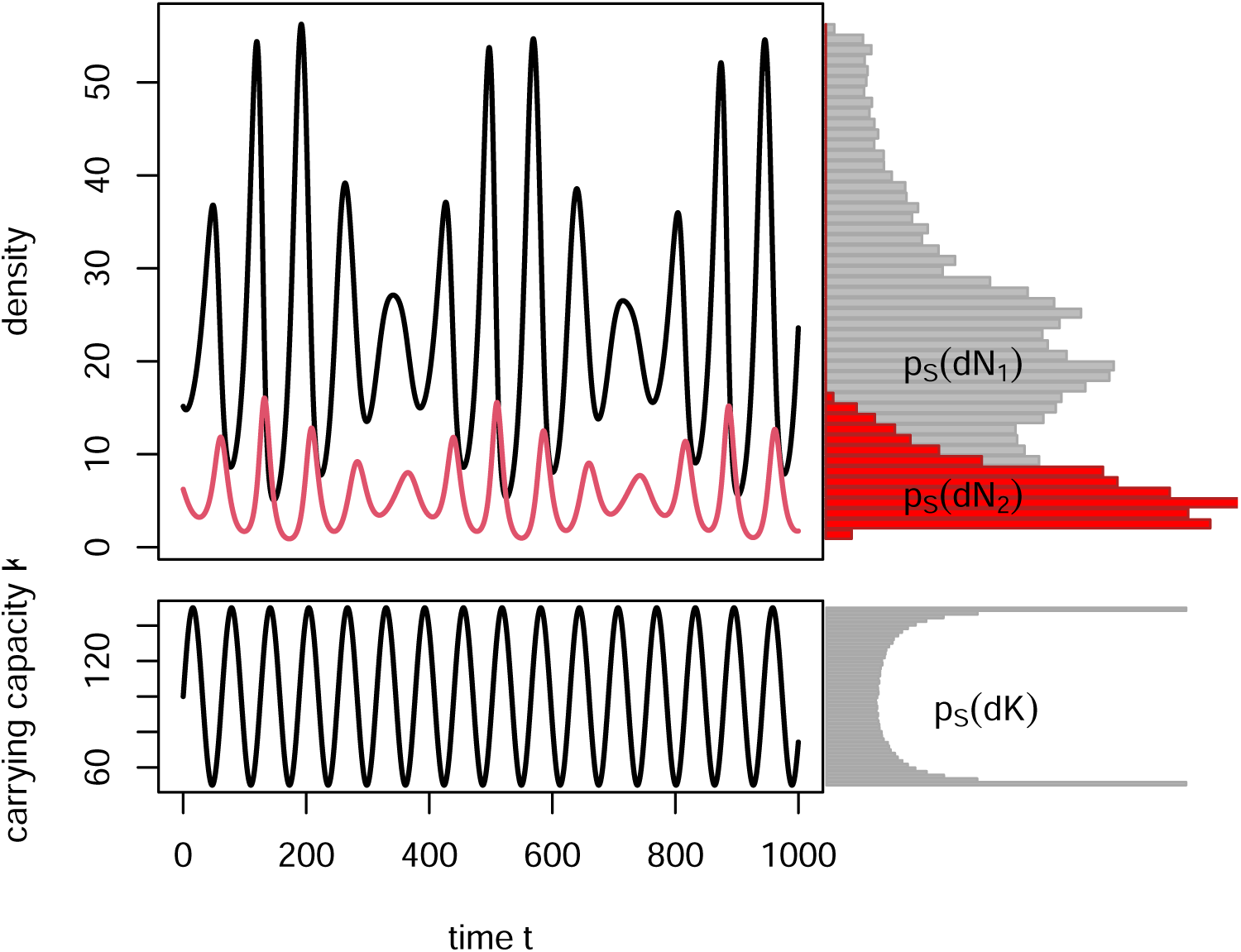
Seasonally forced predator-prey dynamics with marginals *p*(*dN*_1_), *p*(*N*_2_), *p*(*dK*) of the stationary distribution *p*(*dN*_1_, *dN*_2_, *dK*). The (approximate) marginals *p_S_*(*dN*_1_), *p_S_*(*dN*_2_), *p_S_*(*dA*) of this stationary distribution (e.g. *p_S_*(*dN*_1_) = ^∞^ *p_S_*(*dN*_1_, *dN*_2_, *dA*)*dN*_2_*dN*_2_)) are plotted as histograms.

## Appendix E Alternative insights from the invasion graph

**Figure A2:**
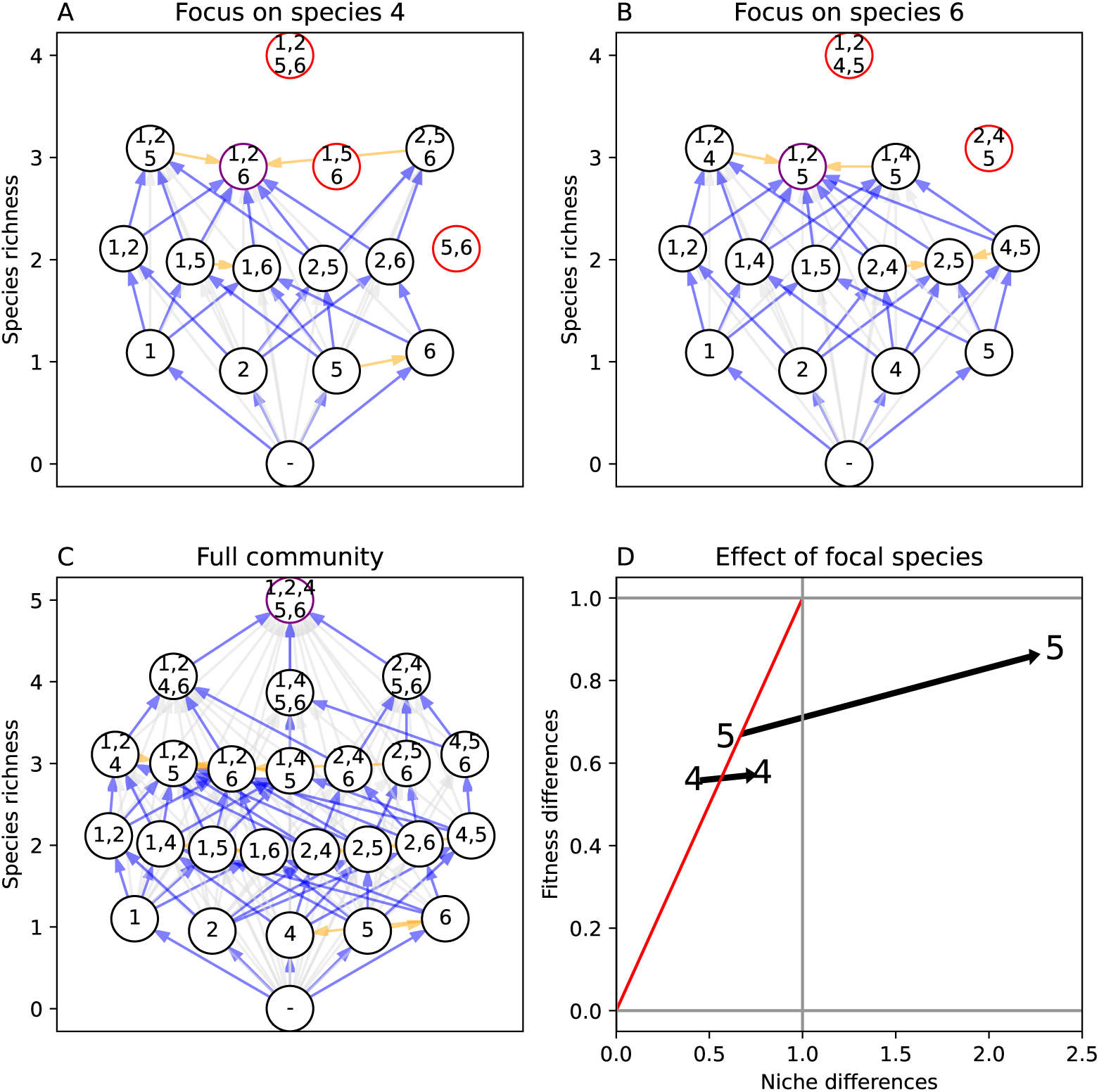
We investigate what hampers the formation of the *n* − 1 communities for species 4 and species 6. We first focus on species 4: Panel A shows invasion graphs for the sub-communities when species 4 is absent. The community 1, 2, 6 is invasion resistant with respect to all species present in panel A, yet species 4 can invade and would lead to community 1, 2, 4, 6 (panel C). Species 5 could not invade into community 1, 2, 6, as its niche differences were not sufficient to overcome it’s fitness differences. Yet, the invasion of species 4 increases the niche differences of species 5 which can no invade (pandel D). For species 6: Similarily, the community 1, 2, 5 is invasion resistant with respect to species 4, but species 6 can invade and would lead to community 1, 2, 6, as species 5 would go extinct. Species 6 increases the niche differences of species 4 without much affecting its fitness differences.

**Figure A3:**
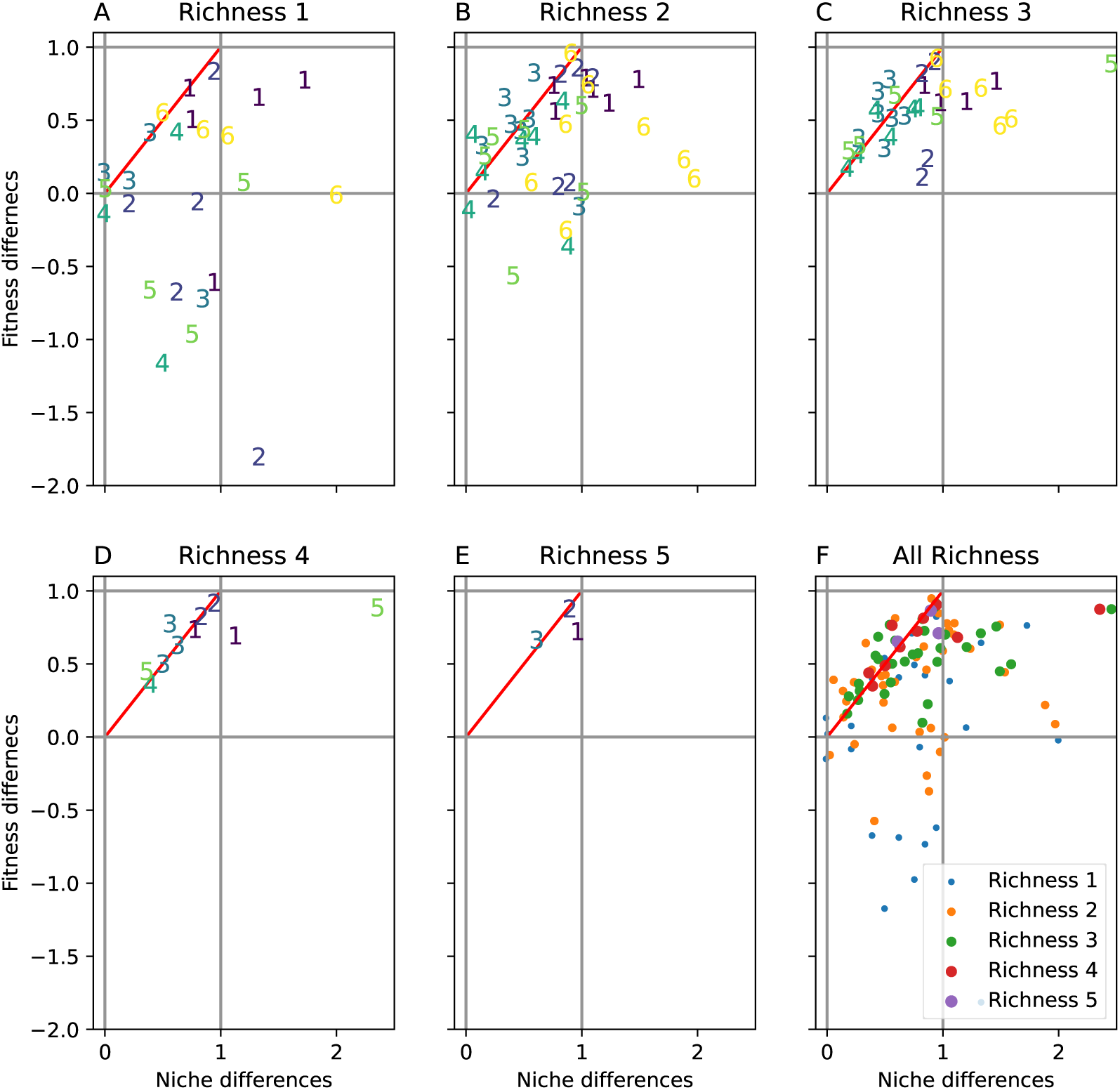
For the community shown in figure 3 we computed niche and fitness differences along the community assembly. Panels A-E show the niche and fitness differences according to the species richness in the resident community. We note that increasing species richness reduces the spread of the niche and fitness differneces and generally increases fitness differneces, which likely limits the species richness. This reduction in spread is not solely due to a reduction in datapoints, specifically panesl B and C have more datapoints than panel A. Panel F shows niche and fitness differences of all richness. Panel A is equivalent to Figure 3 B. However, the datapoints from Figure 3 D are split into palen D and panel E, as species 3 invades into a community with 5 species. On the other hand, panels D and E contain datapoints not present in Figure 3D, as we focus on all possible communities with that richness, not only on a subset.

